# Pinpointing the tumor-specific T-cells via TCR clusters

**DOI:** 10.1101/2022.01.30.478375

**Authors:** M.M. Goncharov, E.A. Bryushkova, N.I. Sharayev, V.D. Skatova, A.M. Baryshnikova, G.V. Sharonov, I.V. Samoylenko, L.V. Demidov, D.M. Chudakov, E.O. Serebrovskaya

## Abstract

Adoptive T cell transfer (ACT) is a promising approach to cancer immunotherapy, but its efficiency fundamentally depends on the extent of tumor-specific T-cell enrichment within the graft. This can be estimated via activation with identifiable neoantigens, tumor-associated antigens (TAAs), or living or lyzed tumor cells, but these approaches remain laborious, time-consuming, and functionally limited, hampering clinical development of ACT. Here, we demonstrate that homology cluster analysis of T cell receptor (TCR) repertoires efficiently identifies tumor-reactive TCRs allowing to: 1) detect their presence within the pool of tumor-infiltrating lymphocytes (TILs); 2) optimize TIL culturing conditions, with IL-2_low_/IL-21/anti-PD-1 combination showing increased efficiency; 3) investigate surface marker-based enrichment for tumor-targeting T cells in freshly-isolated TILs (enrichment confirmed for CD4^+^ and CD8^+^ PD-1^+^/CD39^+^ subsets), or re-stimulated TILs (informs on enrichment in 4-1BB-sorted cells). We believe that this approach to the rapid assessment of tumor-specific TCR enrichment should accelerate T cell therapy development.

## Introduction

Tumors develop diverse mechanisms of immune evasion, including the generation of hypoxic conditions^1^, inflammation^2^, establishment of an immunosuppressive microenvironment^3–6^, downregulation of antigen presentation^7–9^, promotion of regulatory T cell (T_reg_) infiltration^10^ and outgrowth^11^, and induction of T cell dysfunction^12^. The infusion of large numbers of expanded autologous tumor-reactive T cells—typically after the implementation of lymphodepleting regimens—represents a powerful therapeutic option that may override these immunosuppressive mechanisms. Clinical protocols for such adoptive cell transfer (ACT) therapeutic strategies^13^, inspired by the pioneering work of Steven Rosenberg’s group^14–16^, are now being actively developed and used to treat patients^17–21^.

There is accordingly great demand for methods for the enrichment of autologous tumor antigen-specific T cells for use in ACT protocols. Current techniques rely on the identification of patient-specific peptide neoantigens, which are then used for the functional characterization and selection of cultured tumor-infiltrating lymphocytes (TILs)^22,23^. Alternatively, since the identification of unique neoantigens is costly, time-consuming, and functionally limited in terms of the spectra of identifiable antigens, cultured autologous tumor tissue can be used as a source of antigen-specific stimulus^24^. Certain cell-surface markers such as PD-1, CD39, CD69, CD103, or CD137 may also help to delineate T cell subpopulations (typically CD8^+^) that are enriched for clonally-expanded tumor-reactive T cells^25–27^, therefore culturing selected TIL subsets, such as PD-1^+^ T cells^28^, is a feasible option. In all of these scenarios, however, there is the need for a robust method that enables estimation of enrichment of the transplanted cells with tumor-specific T cells based on the T cell receptor (TCR) repertoire without prior knowledge of TCR specificities.

The ongoing adaptive immune response is often driven by groups of T cell clones with highly homologous TCR sequences that recognize the same epitopes^29–32^. If one can properly account for the most public TCR clusters, which arise regardless of their binding properties by virtue of their high probability of being generated in course of V(D)J recombination^33,34^, identification of such convergent TCR clusters becomes a highly efficient approach to capture clonotypes involved in the current immune response^35^.

Here, we have employed cluster analysis to identify groups of TCR clonotypes involved in the anti-tumor immune response. We demonstrate that this approach successfully pinpoints known TAA-specific TCRs among TIL repertoires in HLA-A*02 melanoma patients. Furthermore, we find that the number of cluster-related clonotypes and the proportion of the bulk TIL repertoire that they occupy grows significantly after anti-PD-1 immunotherapy. We next investigate the TCR content in sorted CD4^+^ and CD8^+^CD39^+^PD1^+^ TILs, and show that these subsets are prominently enriched for TCR clusters, a substantial fraction of which consists of tumor-specific TCRs. These results provide a rationale for focusing on CD39^+^PD1^+^ TILs in adoptive cancer therapy. Finally, we show that repertoire analysis facilitates optimization of TIL culturing conditions, and allows estimation of the extent of tumor-specific T cell enrichment in cultured donor cells and sorted TAA-activated T-cells. Altogether, our findings strongly support the use of cluster TCR analysis as a powerful tool with practical applications in clinical ACT.

## Results

### TIL clusters include TAA-specific TCRs and grow after immunotherapy

We first analyzed published TIL TCR beta chain (TCRβ) repertoires obtained before and after anti-PD-1 immunotherapy for two cohorts comprising 21 and 8 patients with cutaneous melanoma^36,37^. Using the ALICE alrorithm^29^, we were able to identify clusters of convergent TCRβ clonotypes in all patients. The number of cluster-related clonotypes significantly increased after therapy in both cohorts (p = 0.019 and 0.038, respectively; **Fig. 1a,b**), which might reflect treatment-dependent expansion of convergent antigen-specific TILs. A VDJdb database^38,39^ search identified cluster-related clonotypes that are highly similar or identical to known TCRβ variants specific to the HLA-A*02-restricted melanoma-associated antigens Melan-A (Melan-A_aa26-35_ ELAGIGILTV) and NY-ESO-1 (NY-ESO-1_aa157-165_ SLLMWITQC) in 20% and 50% of patients from the Ref. 36 and Ref. 37 cohorts, respectively. Cluster-related clonotypes were enriched for TAA-specific TCRβ variants compared to the bulk TCR repertoire, indicating that cluster analysis can capture clonotypes involved in an ongoing anti-tumor immune response (**Fig. 1c,d**). **Fig. 1d** shows identified TCR clusters for one of the patients after immunotherapy. Summary for the count and size of TCR clusters before and after immunotherapy for each patient is shown on **Fig. 1e,f**.

**Figure 1.**
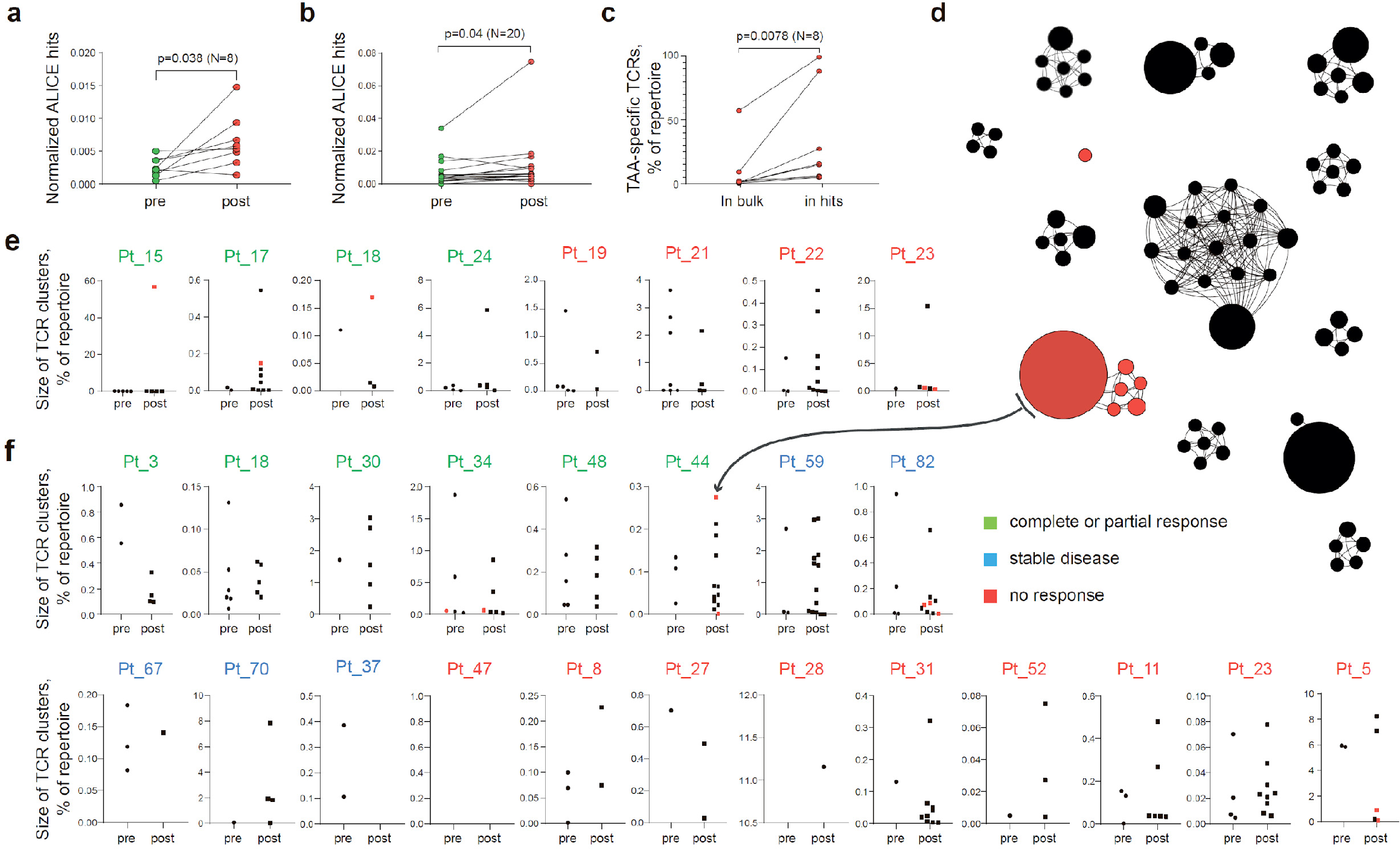
Melanoma TIL TCR clusters before and after immunotherapy. **a,b**. Normalized counts of cluster-related TCRβ clonotypes from metastatic melanoma samples before and after anti-PD-1 therapy from datasets published in (a) Ref. 36 and (b) Ref. 37. **c**. Cumulative frequency of VDJdb-matched TAA-specific clonotypes within the whole repertoire (in bulk) and within cluster-related clonotypes (ALICE hits) of patients with at least one VDJdb-matched cluster (N = 8 patients) from the two published datasets. **d**. TCRβ clusters from patient pt44 (Ref. 36). VDJdb-matched TAA-specific clonotypes are colored in red. **e, f**. Number and size of clusters before and after therapy for each patient from (e) Ref. 36 and (f) Ref. 37, one point corresponds to one cluster. VDJdb-matched TAA-specific clusters are colored in red. N = number of biological replicates in each group. Data in (a) were analyzed with paired t-test; b, c were analyzed with the Wilcoxon test.

Notably, the HLA genotypes of the patients for these two cohorts were unknown. Since VDJdb currently includes limited diversity of HLA contexts, we believe that a much higher proportion of cluster-related clonotypes will be assigned to TAA-specificities with the accumulation of TCR specificity data from more diverse HLA contexts.

### CD39^+^PD1^+^ TILs are enriched with clonal, convergent, and tumor-specific TCRs

It was previously reported that CD39^+^ and PD-1^+^ TIL subsets can be enriched for tumor-specific T cells^25,28,40–45^. To verify whether there is concurrent enrichment with convergent TCR clusters, we performed fluorescence-activated cell sorting (FACS) of CD39^+^PD-1^+^ (double-positive, DP) and non-DP CD4^+^ and CD8^+^ T cells from TILs freshly isolated from lymph node metastases from eight melanoma patients (**Fig. 2a, Fig. S1a**). Overall, TIL composition was skewed towards greater prevalence of CD4^+^ T-cells, although cells with the DP phenotype were more prevalent among CD8^+^ TILs compared to CD4^+^ cells (**Fig. S2a,b**). TCRβ repertoire analysis revealed increased clonality and cluster enrichment for both CD4^+^ and CD8^+^ DP TILs compared to the corresponding non-DP subsets (**Fig. 2b-g**), with greater clonality amongst CD8^+^ TILs than CD4^+^ TILs regardless of immune checkpoint expression status (**Fig. S2c,d**).

**Figure 2.**
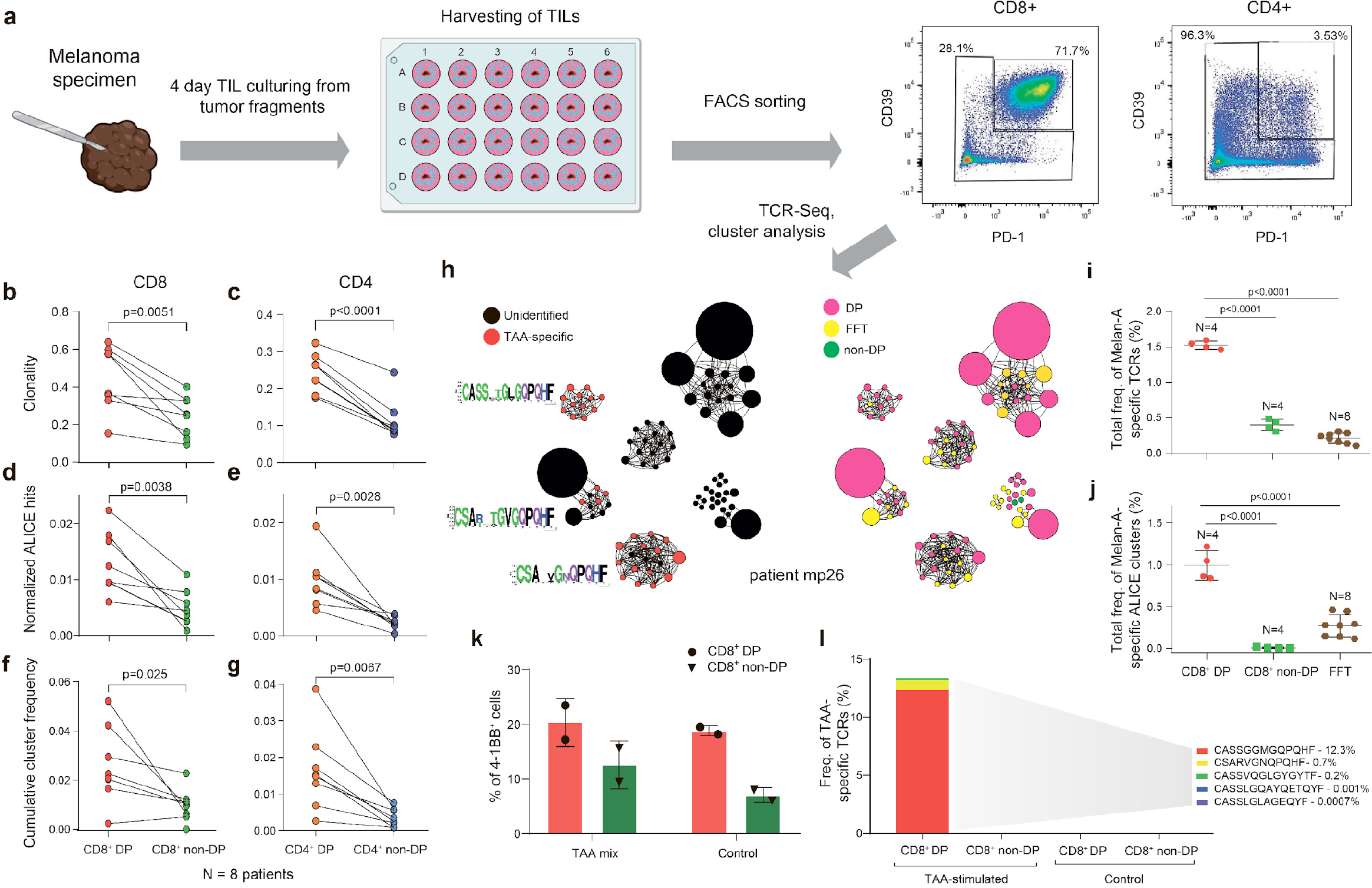
TCR clusters in CD39^+^PD1^+^ TILs. **a**. The experimental workflow. **b**-**g**. TCRβ repertoire analysis for CD8^+^ (b, d, f) and CD4^+^ (c, e, g) DP and non-DP TIL subsets sorted from metastatic lymph nodes of eight melanoma patients. Panels show repertoire clonality calculated as [1-Normalized Shannon-Wiener index] (b, c), normalized counts (d, e) and cumulative frequency of cluster-related clonotypes (f, g). Paired t-test. **h**. TCRβ clusters identified in repertoires obtained from fresh-frozen tumor samples (FFT), and sorted CD8^+^ DP and non-DP TILs for HLA-A*02 patient mp26. VDJdb-matched TAA-specific clusters are colored in red. **i,j**. Cumulative frequency of (i) VDJdb-matched TAA-specific clonotypes and (j) VDJdb-matched TAA-specific cluster-related clonotypes within CD8^+^ DP, CD8^+^ non-DP, and FFT TCRβ repertoires of patient mp26. One-way ANOVA, Bonferroni multiple comparisons correction. **k,l**. Proportion of CD137^+^ cells among CD8^+^ T cells (k) and proportion of VDJdb-matched TAA-specific clonotypes in sorted CD137^+^CD8^+^ T cells (l) in DP and non-DP TILs from patient mp26 that were cultured and re-stimulated with TAA-loaded or control dendritic cells.

For TILs obtained from HLA-A*02-positive patient mp26, with BRAF^wt^ melanoma, a VDJdb search identified three TCR clusters that included A*02-Melan-A_aa26-35_-specific clonotypes (**Fig. 2h**). Both the Melan-A-specific clonotypes and the TCR clusters comprising such clonotypes were prominently enriched within the CD8^+^ DP subset, compared to the bulk TCR repertoire obtained from fresh-frozen tumor tissue (FFT) and to the CD8^+^ non-DP subset (**Fig. 2i,j**). Similar results were obtained for another BRAF^wt^ HLA-A*02-positive patient, pt41 (**Fig. S1b,c**). We concluded that the CD39^+^PD1^+^ fraction is enriched for large and convergent T cell clones that are involved in an ongoing tumor-specific immune response, a substantial portion of which are detectable via cluster TCR analysis.

To functionally confirm our findings, we used the CD137 (4-1BB) upregulation assay. Sorted DP and non-DP TIL subsets from patient mp26 were cultured for two weeks and stimulated with autologous monocyte-derived dendritic cells loaded with a TAA peptide mix. CD8^+^CD137^high^ subsets were subsequently quantified with flow cytometry and sorted for TCRβ library preparation. As shown in **Figure 2k**, the proportion of CD137^high^ cells was higher in cultured DP cells, but we found no difference between T cells stimulated by TAA-loaded or control dendritic cells. At the same time, TCRβ repertoire analysis revealed that the CD137^high^ fraction of TAA-stimulated CD8^+^ DP—but not non-DP or control DP—cells was enriched with known Melan-A-specific clonotypes (**Fig. 2l**). These clonotypes included a TAA-reactive TCRβ variant, CSARVGNQPQHF-TRBV20-TRBJ1-5, which was previously detected in cluster analysis of non-cultured CD8^+^ DP cells, and variant CASSGGMGQPQHF-TRBV19-TRBJ1-5, which is homologous to another cluster. TAA-specific clonotypes cumulatively occupied ~13% of the CD137^high^ fraction of the cultured and TAA-activated DP TILs. These results underscore the importance of TCR repertoire analysis of responding cells, even in the apparent absence of a quantifiable difference between antigen and control conditions, and demonstrate that CD137 marker analysis on its own is insufficiently informative.

### TCR cluster analysis facilitates optimization of TIL culturing conditions

We next investigated the effect of four distinct TIL culture conditions on the expansion of tumor-reactive T cells: IL-2_high_, IL-2_low_/IL-21, IL-2_low_/IL-21/anti-PD-1, and IL-2_low_/IL-21/anti-PD-1/IFNɣ (**Fig. S3a**), where the anti-PD-1 agent employed was nivolumab and the concentration of IL-2 in the low and high conditions was 100 IU/mL and 3,000 IU/mL, respectively. For each condition, we analyzed TCRβ repertoires of TILs independently cultured from 12 tumor fragments collected from patient mp26. TCR clusters identified from all samples were joined and visualized along with the non-cultured DP and non-DP CD8^+^ subsets described above (**Fig. 3a**). As a readout, we used the following:

**Figure 3.**
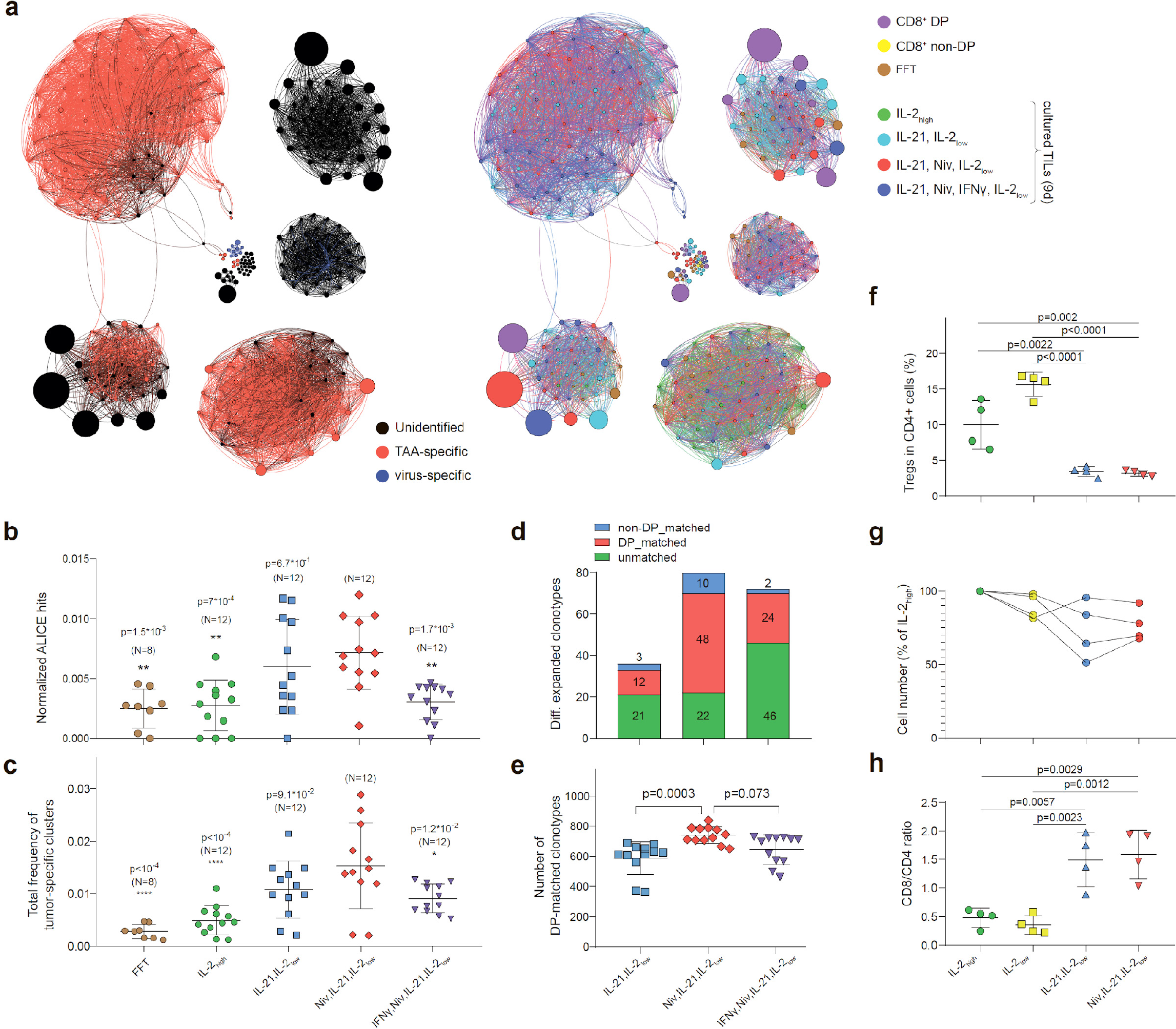
Influence of culture conditions on TILs and TCR clusters. **a**. Combined TCRβ clusters for FFT, CD8^+^ DP, CD8^+^ non-DP, and four different TIL culture conditions on tumor samples from patient mp26. Left: colors indicate VDJdb-defined clonotype specificites. Right: colors indicate the sample of origin. **b–e**. TCRβ repertoire analysis for patient mp26 TILs cultured in four distinct conditions. Panels show (b) normalized counts of cluster-related TCRβ clonotypes, (c) proportion of the repertoire occupied by Melan-A-specific TCRβ clusters, (d) count and subset matching of clonotypes preferentially expanded compared to the IL-2_high_ condition, and (e) the count of CD8^+^ DP matched clonotypes among the top 1,500 clonotypes. N = number of separate tumor fragments. **f**. CD8^+^/CD4^+^ ratio for TILs from patient mp41 in each culture condition. **g**. Fraction of T_reg_s (CD25^+^CD127^-^FoxP3^+^) among CD4^+^ T cells in TILs from patient mp41 in each culture condition. **h**. TIL counts relative to IL-2_high_ culture conditions; n = 4 patients. Data in b,c,f,g were analyzed with one-way ANOVA, Bonferroni multiple comparisons correction, with each condition compared to the IL-21/IL-2_low_/anti-PD-1 condition. Data in the panel (e) was analyzed with Kruskal-Wallis test, Dunn’s multiple comparison test, with each group compared to IL-21/IL-2_low_/anti-PD-1.

i. normalized count of cluster-related clonotypes (**Fig. 3b**),
ii. cumulative proportion of the repertoire occupied by Melan-A-specific TCRβ clusters (clusters predominantly comprising VDJdb-defined Melan-A-specific clonotypes) (**Fig. 3c**), and
iii. the number of differentially-expanded clonotypes compared to pan-activating IL-2_high_ culture conditions, and the proportion of such expanded clonotypes that were also initially detected among CD8^+^ DP TILs (**Fig. 3d**).

The IL-2_low_/IL-21/anti-PD-1 combination yielded the greatest number of cluster-related clonotypes and the highest cumulative proportion of Melan-A-specific clusters out of all culture conditions (**Fig. 3b,c**). Addition of IFN γ did not further enhance the expansion of potentially tumor-specific clones.

We utilized edgeR^46^ software, which was initially designed for differential gene expression analysis, to identify clonotypes that were significantly expanded in tumor fragment cultures in the presence of IL-21 and IL-2_low_ (either with or without nivolumab and IFNɣ) compared to classical pan-activating IL-2_high_ culture conditions. The IL-2_low_/IL-21/anti-PD-1 combination yielded the highest number of reproducibly expanded clonotypes, 60% of which were detected among freshly-sorted CD8^+^ DP TILs (**Fig. 3d, Fig. S3b**). The overall count of CD8^+^ DP-matched clonotypes was also highest for this combination (**Fig. 3e**). These results show the positive influence of PD-1 inhibition on the proliferative potential of CD8^+^CD39^+^PD-1^+^ T cells. We also noted a slight increase in the number of CD8^+^ non-DP matched clonotypes which were expanded in these same conditions—10 clonotypes, compared to three clonotypes in IL-2_low_/IL-21 without nivolumab (**Fig. 3d**)—which can be explained by nivolumab-dependent expansion of PD-1^+^CD39^-^ T cells.

### IL-2_low_/IL-21/anti-PD-1 culture conditions stimulate TIL proliferation without T_reg_ expansion

T_reg_s are known to hamper anti-cancer immune responses elicited by *ex vivo* expanded TILs^47^. This immunosuppression may be overcome by CD25^+^ T cell depletion^48^. Alternatively, T_reg_ expansion may be suppressed by IL-21^49^. To more comprehensively characterize the distinct TIL culture conditions described above, we compared their capacity to support or suppress T_reg_ expansion and overall T cell expansion. Predictably, IL-2 alone yielded the highest proportion of T_reg_ (CD4^+^CD25^high^CD127^low^FoxP3^high^ cells) cells. In line with prior findings^50^, IL-2_low_ created preferential conditions for selective T_reg_ expansion, while the addition of IL-21 drastically reduced the proportion of T_reg_s amongst CD4^+^ TILs (**Fig. 3f, Fig.S3c,d**).

IL-21 was previously demonstrated to promote the expansion of T cells with a memory phenotype^51^. We also noted a significant increase in the number of memory phenotype^52^ CD127^high^ cells among CD4^+^ lymphocytes in IL-21^+^ conditions regardless of the presence or absence of nivolumab (**Fig. S3e**). Remarkably, CD8^+^ TILs displayed significant CD127^high^ enrichment only upon simultaneous introduction of IL-21 and nivolumab (**Fig. S3f**). These results reveal the synergistic action of IL-21 and disruption of PD-1-dependent signaling by nivolumab on expansion of CD8^+^ memory T cells.

Regarding overall proliferative potential, the highest T cell count was evident for IL-2_high_ conditions, although we also observed roughly comparable numbers in the IL-2_low_ cultures (**Fig. 3g**). The presence of IL-21 stifled IL-2-dependent TIL proliferation, as previously reported for human CD4^+^ T cells^53^. On the other hand, IL-21 was favorable for expansion of CD8^+^ TILs, whereas IL-2 alone favored CD4^+^ cell growth (**Fig. 3h**).

## Discussion

Here we demonstrate that rational TCR clustering can be used to identify tumor-specific T cell clones and estimate their relative enrichment among TILs. Starting with published TCR repertoire data from melanoma tumors, we identified clusters of convergent TCR clonotypes, which increased in numbers and total frequency after anti-PD-1 immunotherapy. We next found significant enrichment of convergent TCR clusters in PD1^+^CD39^+^ subpopulations of both CD4^+^ and CD8^+^ TILs, which were previously shown to be enriched in terms of tumor reactivity^41,43,54^. These findings are further supported by the data we obtained for cases where we know the HLA context, as well as some TCR clonotypes of interest and their cognate antigens. A VDJdb database search successfully identified TCR variants specific to TAA antigens in HLA-A*02-positive patients, where approximately half of the clusters could be matched to known Melan-A-specific sequences. The cumulative frequency of such Melan-A specific clusters within the CD8^+^ DP population was only slightly lower than the total frequency of all identified Melan-A specific TCRs, indicating that our approach was able to identify most of the high-frequency tumor-specific clonotypes. It should be noted that TCR clusters are an essential feature of a convergent immune response that involves several homologous clonotypes. For those cases where a single T cell clone dominates in response to a particular antigen and/or homologous “neighbours” are absent due to the very low probability of convergent TCR generation^30^, cluster analysis may miss some of the tumor-reactive TCRs. Nevertheless, our results demonstrate the overall power of this approach, which is applicable in those situations where specific antigens are unknown.

We describe one potential implementation of our approach by using it to optimize conditions for *ex vivo* TIL expansion. In particular, TILs cultured in IL-21^+^ conditions demonstrated the highest number of cluster-related TCRs, which is indicative of a more prominent influence of antigen-driven TCR selection. We speculate that IL-21 exerts its influence in our culture system both at the antigen-presentation stage (as MHCI- and, to a lesser extent, MHC II-restricted antigen presentation by tumor cells occurs while tumor fragments are cultured *ex vivo*)^55^ and at the expansion stage, where IL-21 selectively promotes the expansion of non-regulatory T cell populations. The consequences of *in vitro* PD-1 blockade are less well understood. The *in vitro* application of an anti-PD1 antibody was previously shown to induce IFNɣ and TNFα secretion by PD1-expressing CD8^+^ TILs^56^. In our work, we found a greater increase in abundance of cluster-related and tumor-specific TCRs for cells cultured in IL-21^+^/anti-PD-1 conditions compared to IL-21^+^ alone.

We expect that this straightforward TCR repertoire-based approach to estimation of TIL enrichment with tumor-reactive clones will accelerate clinical development of adoptive T cell therapy. At the R&D stage, this approach could greatly facilitate the selection of optimal TIL subsets and optimization of TIL culturing conditions and downstream enrichment procedures. And within clinical pipelines, such analyses should make it possible to estimate tumor-reactive TIL abundance at the level of individual tumor samples, both before and after culturing and/or enrichment. As the field of T cell therapy advances towards the ability to determine the clonal specificities and phenotypes that can effectively fight each individual patient’s tumor, methods for grouping convergent TCRs that respond to the same tumor antigens will become an essential part of the toolbox for rationally-designed T cell therapy.

## Acknowledgements

We thank the Center for Precision Genome Editing and Genetic Technologies for Biomedicine (Moscow) for the genetic research methods, and personally V. Cheranev and D. Korostin for HLA typing of patient tumor material. We thank Valeria V. Kriukova for the help with data analysis. We thank Skoltech Genomics Core Facility for performing high-throughput sequencing of mp41 libraries (Skoltech Life Sciences Program grant for the use of shared facilities, received by M.Goncharov). We are grateful to Michael Eisenstein for his valuable help in editing the manuscript.

## Online methods

### Patients

All clinical samples were acquired from the N.N. Blokhin National Medical Research Center of Oncology in accordance with protocol MoleMed-0921, approved by the ethical committee on 30 Jan 2020. All patients involved in the study were diagnosed with metastatic melanoma and signed an informed consent prior to collection of their biomaterial. Most experiments were performed on freshly resected metastatic lymph nodes obtained during surgery. Genotyping for BRAF^V600E^ was performed at the N.N. Blokhin National Medical Research Center of Oncology, Pathomorphology Department. From each patient enrolled in this study, we also obtained 20–30 mL of peripheral blood before the surgery. Patient information is provided in **Table S1**.

### Brief TIL culture

Freshly-resected tumor specimens were dissected into fragments measuring 1–3 mm in each dimension. Several fragments were frozen in liquid nitrogen for further cDNA library preparation and HLA-typing. Individual fragments were seeded into the wells of a 24-well tissue culture plate with 2 mL of complete T cell cultivation medium (CM) supplemented with 5% heat-inactivated human AB serum (PanBiotech, Germany) and 1,000 IU/mL IL-2 (Ronkoleukine, BiotechSpb, Russia) CM consisted of RPMI-1640 (PanEco, Russia), 25 µM/mL HEPES, pH 7.2 (PanEco, Russia), 100 IU/mL penicillin, 100 µg/mL streptomycin, 10 µg/mL gentamicin, 1x non-essential amino acids mix, 1x GlutaMAX β-mercaptoethanol (0.55 µM) (all from Gibco, Thermo Fisher Scientific, US) and 110 μg/m sodium pyruvate. On the fourth day of cultivation, TILs were harvested, filtered through a 70-μm mesh, stained with fluorophore-labeled antibodies, and FACS sorted. For the generation of TCR repertoire libraries, T cells were sorted directly into the RLT cell lysis buffer (QIAGEN, Netherlands) and stored at -80ºC until used for RNA isolation. For functional assays, TILs were sorted into 1.5 mL Eppendorf tubes with 0.5 mL RPMI-1640. Live-sorted T cells were seeded into wells of 96-well cell culture plate at 10^6^ cells/mL in CM and cultured for at least five days. Here, CM was supplemented with 1,000 IU/mL IL-2, 50 ng/mL IL-21 (SCI-STORE, Russia), 20 μg/mL nivolumab (Bristol-Myers Squibb, USA), and 10% autologous patient-derived serum. Half of the media was replaced three times a week. One day before the functional assays, all the media was replaced with fresh CM without interleukins or nivolumab.

### Expansion of bulk TILs

Fragments of tumor specimens from each of four melanoma patients (mp26, 32, 34, and 42) were plated into separate wells of a 24-well tissue culture plate (1 fragment per well, 1 plate per patient) with basic RPMI-1640-based cultivation medium and 5% heat-inactivated human AB serum. Bulk TILs were expanded from tumor fragments in the following different conditions (6 wells/condition) for 9 days: 1) IL-2_high_ (3,000 IU/mL), 2) IL-2_low_ (100 IU/mL), 3) IL-21 (25 ng/ml) + IL-2_low_, 4) nivolumab (20 μg/ml) + IL-21 + IL-2^low^, and 5) IFNγ (100 μg/ml) + nivolumab+ IL-21+ IL-2_low_. TILs from patient mp26 were cultivated in conditions 1, 3, 4 and 5; TILs from patients mp32, 34, and 42 were cultured in conditions 1–4. Half of the medium was replaced on days 3 and 5 with the same medium as was used for initial plating. On day 9 of cultivation, TILs from each well were counted and lysed in RLT buffer at 5 × 10^5^ cells/mL density for preparation of TCR repertoire libraries.

### Expansion of sorted T-cells

FACS-sorted T-cells were expanded using non-specific TCR-dependent stimulation with anti-CD3/CD28 Dynabeads (Thermofisher Scientific, USA). Beads were added into the cultivation medium on the next day after seeding the cells, with 2 µl of bead solution per 10^5^ cells. Sorted TILs were expanded for 2 weeks. Beads were magnetically removed upon achieving desired cell numbers. Before co-cultivation experiments, cells were allowed to “rest” for one day in interleukin-free CM supplemented with 10% autologous patient-derived serum (*i*.*e*., “resting” medium).

### PBMC isolation

Peripheral blood mononuclear cells (PBMCs) were derived from patients’ blood samples using gradient centrifugation with Ficoll-Paque Plus (GE Healthcare). Briefly, 18 mL of whole blood were diluted to 50 mL volume with sterile 1x PBS. Diluted blood was layered over the Ficoll-Paque solution in 50 mL SepMate tubes (StemCell Technologies, USA), with 25 mL of diluted blood per tube. SepMate tubes were centrifuged for 20 min at 1200 x *g* with brake off. Afterward, buffy coats were collected and washed two times with 50 mL of sterile PBS.

### Monocyte-derived dendritic cells cultivation

Autologous dendritic cells were generated as described in Ref. 57. Briefly, CD14^+^ cells (monocytes) were isolated from patients’ PBMCs with a magnetic enrichment procedure using anti-CD14 MicroBeads (Miltenyi Biotec, Germany). Then, monocytes were seeded into the wells of 24-well tissue culture plates at 5 × 10^5^ cells/well. X-Vivo-15 medium (Lonza, Switzerland) with 400 U/mL IL-4 and 800 U/mL GM-CSF was used for the differentiation of monocytes. On the fourth day of cultivation, the medium was renewed, and dendritic cells were loaded with the mix of melanoma TAA peptides (PepTivator Melan-A/MART-1, gp100/Pmel, and MAGE-A3 human; Miltenyi Biotec) at a concentration of 600 nM each. The next day, loaded DCs were matured using 1 µg/mL PGE, 10 ng/mL IL-1β, and 25 ng/mL TNF-α. Following 24 hours of maturation, DCs were harvested and used for co-cultivation with T cells.

### CD137 antigen-specific activation assay

FACS-sorted PD1^+^CD39^+^ (DP) and non-DP populations after expansion and two days “rest” without IL-2 were co-cultured with antigen-loaded autologous DCs at a ratio of 10:1 T cells:DCs. The co-cultivation medium consisted of 1:1 CM plus AIM-V serum-free medium (Gibco, USA) supplemented with 50 ng/mL IL-21. Both CD137^high^ and CD137^low^ T-cells were lysed with RLT buffer for further RNA isolation and TCR library construction. The frequency of CD137^high^ cells was measured for both CD4^+^ and CD8^+^ TILs. Antigen-specific activation was measured as a ratio of CD137^high^ T cell frequencies in TILs co-cultured with antigen-loaded vs unloaded DCs.

### Flow cytometry

For FACS sorting of PD1^+^CD39^+^ TIL subpopulations, cells were stained with the following antibodies: CD4-BV510 (RPA-T4), CD8-Alexa-647 (SK1), CD127-APC-Cy7 (IL-7Rα, A019D5), PD-1-BV421 (CD279, EH12.2H7) (BioLegend, Germany), CD25-PE (IL-2Rα, Beckman Coulter), CD39-FITC (eBioA1) (eBioscience, US). Briefly, cells were pelleted, resuspended in the staining solution with fluorescent antibodies, and incubated for 1 h at 4 ºC in the dark. Next, the cells were washed and resuspended in PBS at an approximate density of 5 × 10^6^ cells/mL. T cells of interest were sorted on a FACS Aria III (BD Biosciences, USA). Data analysis was performed with FlowJo software (FlowJo, LLC). To avoid antigen-specific T_reg_-dependent T cell suppression in further functional assays, we identified T_reg_s as CD4^+^CD25^+^CD127^-^ cells^58^ and sorted them separately. For functional assays, we sorted T cells into CD39^+^ PD-1^+^ double-positive (DP) and non-DP (CD39 or PD-1 single-positive and double-negative) subpopulations, with CD4^+^ and CD8^+^ cells together. For TCR library construction, CD4^+^ DP and CD8^+^ DP cells, as well as corresponding non-DP populations, were sorted separately in order to evaluate individual properties of their TCR repertoires. For the CD137 activation assay, T cells were stained with fluorescently-labeled CD4-BV510 (RPA-T4), CD8-Alexa-647 (SK1), CD137-PE (4B4-1) (BioLegend, Germany) antibodies.

### Nuclear staining for FoxP3

First, T cells were stained for surface markers with CD4-BV510 (RPA-T4), CD25-APC (BC96), CD127-Alexa488 (A019D5) (BioLegend, Germany) for 30 min at 4 ºC in the dark. TILs from patient mp41 were also stained with CD8-BV421 (SK1, BioLegend, Germany) to estimate the CD8/CD4 ratio in various cultivation conditions. Next, the cells were washed and resuspended and processed with the eBioscience Intracellular Fixation & Permeabilization Buffer Set according to manufacturer’s instructions with minor alterations. Briefly, stained cells were fixed with Fixation Reagent for 45 min at 4ºC in the dark. Then, the fixed cells were washed three times in 1 mL of Permeabilization Reagent, resuspended in 50 µL of Permeabilization Reagent and blocked for 15 min in 2% human serum. Next, cells were stained with 2 µL of anti-FoxP3-PE (236A/E7) (BD Biosciences, USA) for 1 h in Permeabilization buffer at 4 ºC in the dark, washed, and analyzed with a Navios flow cytometer (Beckman Coulter, US). Finally, the fraction of T_reg_s (CD4^+^ CD25^high^ CD127^low^ FoxP3^high^) out of total CD4^+^ T cells was determined.

### HLA-typing

For patients mp39, mp41, mp42, and mp44, we have only checked for the presence of HLA-A*02. Aliquots of PBMCs or TILs from these patients were stained with anti-HLA-A*02-PE (BD7.2) (BD Biosciences, USA) antibody and analyzed with a Navios flow cytometer. Other patients (**Table S1**) were HLA-typed using NGS at the Center for Precision Genome Editing and Genetic Technologies for Biomedicine (Moscow).

### RNA isolation and TCR library preparation

RNA from fresh-frozen tumor fragments was isolated using TRIzol reagent(Invitrogen). RNA from RLT-lysed cells was extracted using the RNeasy mini kit (Qiagen) according to the manufacturer’s protocol. RNA concentration was measured with the Qubit RNA HS Assay Kit (Thermo Fisher Scientific, US). No more than 500 ng of total RNA was used for cDNA synthesis.

cDNA libraries were generated using the Human RNA TCR Multiplex kit (MiLaboratories), according to the manufacturer’s protocol. We aimed to achieve coverage of 20 paired-end reads per cell for sorted and cultivated TIL populations, and approximately 2 × 10^6^ reads per tumor sample fragment. Sequencing was performed using the Illumina NextSeq platform (2 х 150 bp read length).

### Analysis of TCR repertoires sequencing data

TCRseq data was analyzed with MiXCR software (MiLaboratories) in order to extract TCRβ CDR3 clonotypes. VDJtools software was used for processing of MiXCR output TCR repertoire data, calculation of TCR repertoire diversity, and pre-processing of TCR repertoires (*i*.*e*., pooling of joined TCR repertoires and down-sampling of TCR repertoires)^59^.

### Cluster analysis of TIL TCRβ repertoires

This analysis was performed using the ALICE algorithm^29^. We selected clonotypes with read count >1 to exclude undercorrected erroneous TCR variants that could potentially create false neighbors of abundant clonotypes, distorting the ALICE hit identification process. P_gen_ of amino acid sequences was estimated using Monte Carlo simulation. For each VJ pair, 5 million TCR sequences were simulated, and 20 iterations of the algorithm were performed. Only clonotypes with Benjamini-Hochberg (BH)-adjusted *p <* 0.001 were selected as significant ALICE hits. As the number of ALICE hits strongly depends on the initial variability of the TCR repertoire, we normalized the number of hits between samples based on the number of the top-frequency input clonotypes.

To visualize the resultant clusters of convergently selected TCRs we used the igraph function^60^. This function utilizes the de Bruijn graph method to calculate the distance between amino acid sequences of CDR3 regions. It creates a graph file in GML format where each node represents an individual TCR clonotype and the distance between nodes is proportional to the difference between CDR3 amino acid sequences. Edges connected nodes representing TCR clonotypes with Hamming distance ≤2. Graphs were visualized using Gephi 0.9.2 network analysis platform^61^. The size of the node represents the frequency of the corresponding TCR clonotype. Upon construction of composite graphs including clonotypes from multiple TCR repertoires, identical clonotypes from different TCR repertoires were displayed by separate nodes.

### Matching cluster-related TCR clonotypes to VDJdb

We annotated TCR repertoire data using the VDJdb database of T cell receptors with known specificity^38^. We assumed that TCRs of interest had the same specificity as TCRs from the database if: i) CDR3 regions of compared TCRs differed no more than by one central amino acid substitution, ii) substituted amino acids belonged to the same group based on their R properties (polar, aliphatic, aromatic, positively/negatively charged), and iii) HLA-restriction of TCR clonotypes from the database matched one of the patient’s HLA alleles, if known.

TCR clusters consisting predominantly of TAA-specific clonotypes, but not clonotypes of other specificities, were considered TAA-specific as a whole. VDJdb-unmatched members of the TAA-specific clusters were deemed to possess the same specificity as the whole cluster based on structural similarity to VDJdb-matched clonotypes and were included in the subsequent analysis.

### Analysis of differentially expanded clonotypes with edgeR software

We used a statistical approach implemented in the edgeR package to identify TCRβ clonotypes that were significantly expanded in bulk TILs of patients mp26 and mp34 as described in Ref. 60. Previously, this approach was used to track vaccine-responding clonotype expansion in time. We implemented edgeR for comparison of TCR repertoires of TILs cultivated in experimental conditions 2-4 (described above) and TILs expanded in IL-2_high_ conditions. Six and four biological replicate samples of each cultivation setting were used for the analysis of TCR repertoires from mp26 and mp34, respectively. TCR clonotypes were deemed expanded if the false discovery rate adjusted p value was <0.01 and the log_2_ fold-change was >1.

### Statistical analysis

Statistical analysis was performed using Graph Pad Prism 8.0 (GraphPad Software Inc., USA). All data was reported as mean ± SD. The Shapiro-Wilk test was used for normality estimation in all cases. Names of statistical tests and numbers of biological replicas in each comparison group are provided in the figure legends.

## Data availability

Extracterd TCR repertoires are available at: https://figshare.com/projects/Pinpointing_the_tumor-specific_T-cells_via_TCR_clusters/125284

---

**Supplementary Table S1.** Patients and clinical characteristics.

| patient ID | gender | age | BRAFmut | HLA-A*02 | diagnosis |
| --- | --- | --- | --- | --- | --- |
| mp24 | F | 53 | V600E |  | melanoma |
| mp26 | F | 67 | wt | + | melanoma |
| mp32 | F | 65 | V600E |  | melanoma |
| mp34 | F | 55 | V600E |  | melanoma |
| mp35 | F | 54 | V600E |  | melanoma |
| mp36 | F | 61 | V600E |  | melanoma |
| mp39 | F | 61 | wt |  | melanoma |
| mp41 | F | 57 | wt | + | melanoma |
| mp42 | F | 75 | N/A |  | melanoma |
| mp44 | F | 62 | wt | + | melanoma |

**Supplementary Figure S1.**
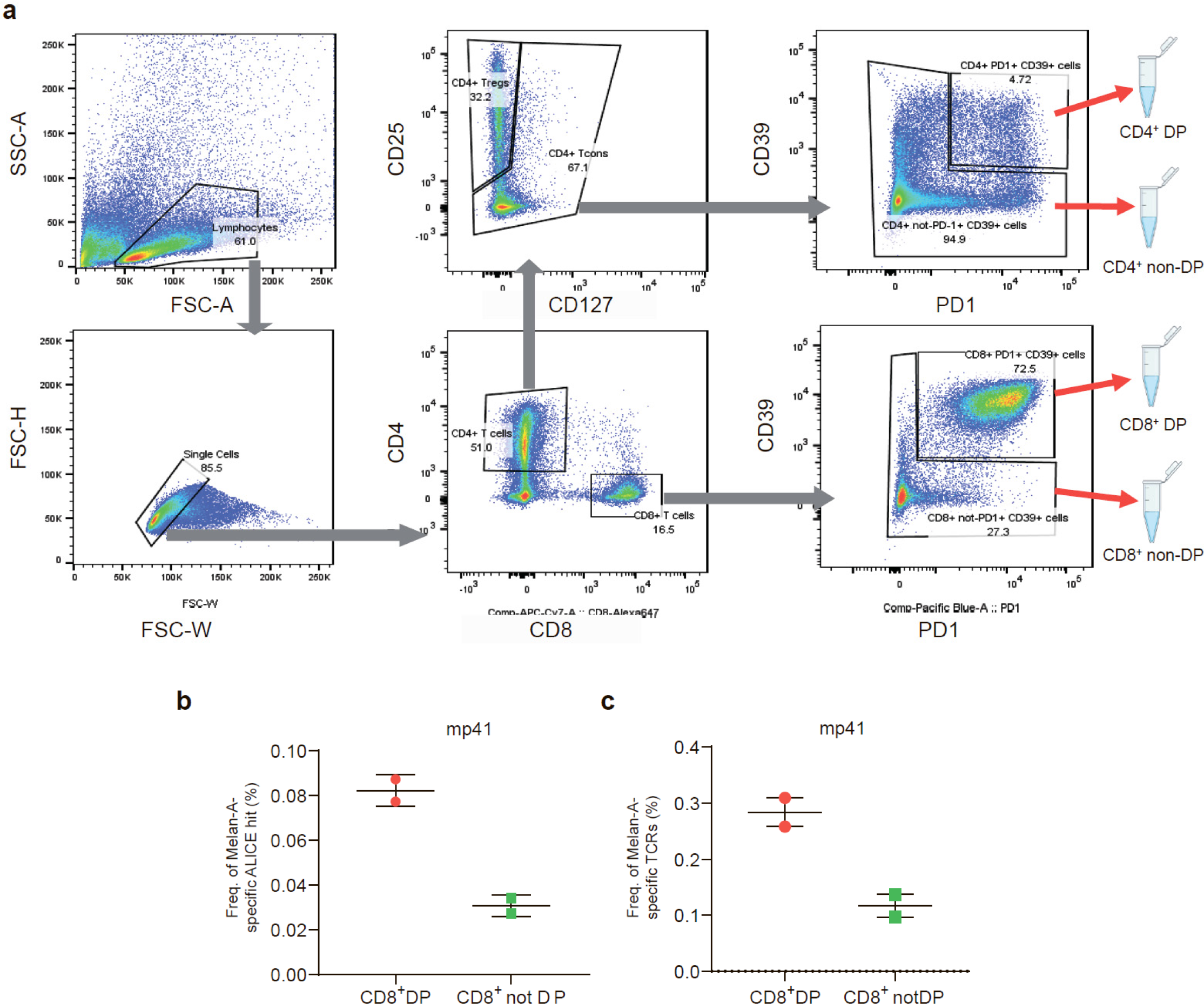
**a**. FACS gating for sorting of CD8^+^ and CD4^+^ CD39^+^PD-1^+^ double-positive (DP) and non-DP T cells from briefly cultured melanoma TILs and single-cell suspensions prepared from tumor samples. **b**. Cumulative frequency of Melan-A-specific cluster-related TCRs in repertoires of sorted CD8^+^ TILs from patient mp41. **c**. Cumulative frequency of Melan-A-specific TCRs in total repertoires of sorted CD8^+^ TILs from patient mp41.

**Supplementary Figure S2.**
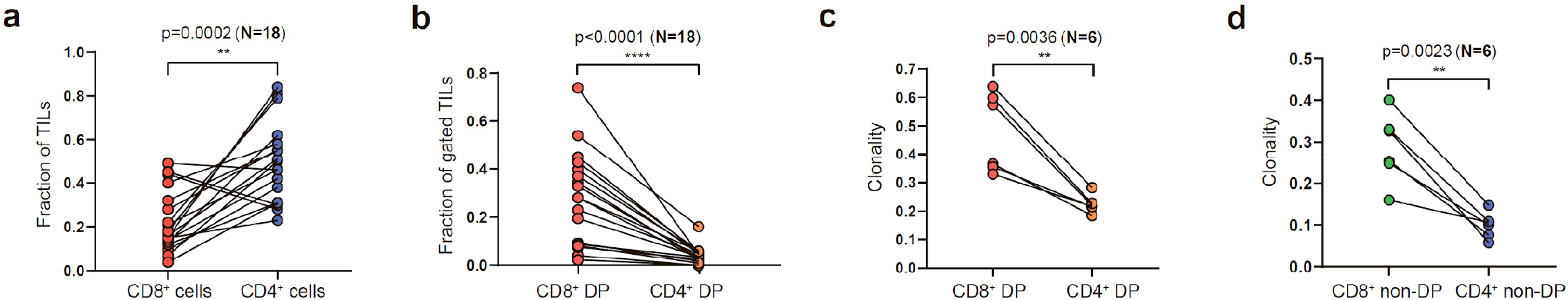
Characteristics of sorted TILs. **a**. Fraction of CD8^+^ and CD4^+^ cells out of all tumor-infiltrating CD3+ T-cells. **b**. Fraction of CD39^+^PD-1^+^ cells consisting of CD8^+^ and CD4^+^ TILs. **c,d**. Clonality (calculated as [1-Normalized Shannon-Wiener index]) of CD8^+^ and CD4^+^ (c) DP and (d) non-DP TILs. N = number of tested patients. Data analyzed by paired t-test.

**Supplementary Figure S3.**
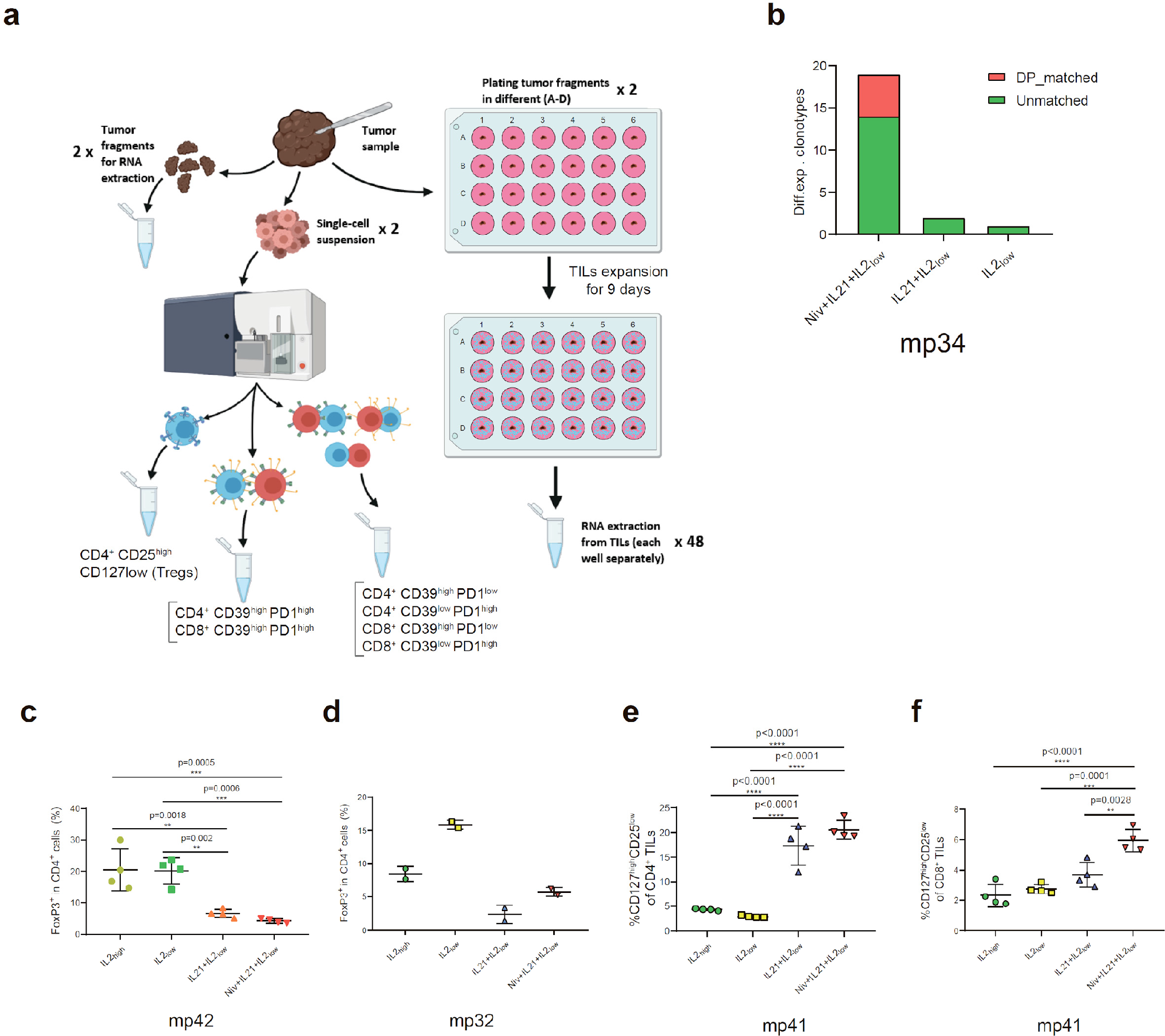
Expansion of TIL subtypes in different cultivation conditions. **a**. Experimental pipeline for evaluating CD39^+^PD-1^+^ T-cell clonotypes expansion from bulk melanoma TILs cultivated in different conditions. **b**. Number of TCR clonotypes preferentially expanded in bulk TILs of patient mp34 under various culture conditions compared to classical IL-2_high_ conditions. **c,d**. Fraction of T_reg_s (gated as CD4^+^CD25^+^CD127^-^FoxP3^+^) among CD4^+^ T cells of patients (c) mp42 and (d) mp32. **e,f**. Proportion of CD127^high^CD25^low^ cells among (e) CD4^+^ and (f) CD8^+^ TIL subsets from patient mp41. Data in c,e,f analyzed by one-way ANOVA, Bonferroni’s multiple comparisons test.

